# Genomic style: yet another deep-learning approach to characterize bacterial genome sequences

**DOI:** 10.1101/2021.08.09.455756

**Authors:** Yuka Yoshimura, Akifumi Hamada, Yohann Augey, Manato Akiyama, Yasubumi Sakakibara

## Abstract

**Motivation:** Biological sequence classification is the most fundamental task in bioinformatics analysis. For example, in metagenome analysis, binning is a typical type of DNA sequence classification. In order to classify sequences, it is necessary to define sequence features. The *k*-mer frequency, base composition, and alignment-based metrics are commonly used. In contrast, in the field of image recognition using machine learning, image classification is broadly divided into those based on shape and those based on style. A style matrix was introduced as a method of expressing the style of an image (e.g., color usage and texture).

**Results:** We propose a novel sequence feature, called genomic style, inspired by image classification approaches, for classifying and clustering DNA sequences. As with the style of images, the DNA sequence is considered to have a genomic style unique to the bacterial species, and the style matrix concept is applied to the DNA sequence. Our main aim is to introduce the genomics style as yet another basic sequence feature for metagenome binning problem in replace of the most commonly used sequence feature *k*-mer frequency. Performance evaluations show that our method using style matrix achieves the superior accuracy than state-of-the-art binning tools based on *k*-mer frequency.

## Introduction

Classification (or clustering) of biological sequences (DNA, RNA, and protein sequence) is one of the most fundamental tasks in bioinformatics analysis. In order to classify sequences, it is necessary to define the sequence features. Three main methods are most frequently used: homology, *k*-mer (word of length *k*) frequency, and base composition (e.g., GC content). In the first method, the homology is scored by performing alignment and calculating the score of the matched positions and the mismatched positions and is then used for classification (Durbin et al., 1998). The *k*-mer frequency, in the second method, is the frequency of occurrence of a sub-sequence (word) of length *k*. A profile is obtained from this frequency distribution, which is expected to be specific to bacterial species, and classification is performed by comparing the similarities of profiles. In the third method, base composition, particularly, the GC content is the percentage of GC among the four bases in the sequence. *S. coelicolor*, a species of actinomycetes that is considered to have a high GC-content, has a large difference in the GC content from 72.2% to 50.6% in *E. coli*. The third method uses this difference in the GC content to classify sequences. Sequences can be classified using these values that characterizes DNA sequences, but there are problems such as the classification accuracy is not always high.

An example of the application of DNA sequence classification is binning, which classifies the obtained DNA sequences to each species in metagenomic analysis (Mande et al., 2012). Metagenome analysis is a method for comprehensively analyzing the DNA extracted from an environment in which multiple species are mixed. Since intestinal flora is known to affect human health and disease, metagenome analysis is attracting attention as a research method to clarify which bacteria are present in the intestinal flora and how they interact with the host. In metagenomic analysis, the total DNA of a mixed sample of various bacteria is extracted and sequenced. The reads obtained by sequencing are subjected to processes such as assembly for reconstructing the genome and binning for classifying the sequences according to the bacterial species. Since the sample contains many unknown bacteria, whose genomes have not been determined, the binning processing that classifies the sequences to each species is important. Most existing binning methods have been developed to bin contigs assembled from sequence reads, and are categorized into supervised (classification) and unsupervised (clustering) approaches. Those methods are further classified into alignment-based and alignment-free ones. The supervised approaches including MEGAN (Huson et al., 2007) and Kraken (Wood and Salzberg, 2014) require the reference database or training data with the taxonomic class labels in order to assign assembled contigs to correct taxonomic classes. In comparison, the unsupervised approaches do not require any additional reference database or taxonomic information. In metagenomic analysis, since many bacteria species in the bacterial flora are unknown and recent next-generation sequencing (NGS) technique generates a large amount of sequence data, the unsupervised and alignment-free methods are more demanded. In the alignment-free method, most binning methods make use of *k*-mer frequency (*k*-mer composition) as the fundamental sequence feature. For example, tetra-mer composition is often used (Chatterji et al., 2008; Yang et al., 2010). In addition, the accuracy is improved by incorporating the coverage information that represents the species abundance in the bacterial flora (MateBAT2 (Kang et al., 2019), CONCOCT (Alneberg et al., 2014), MaxBin2 (Wu et al., 2015), MetaProb (Girotto et al, 2016), and MrGBP (Kouchaki et al, 2019)). However, the accuracy of binning that relies on *k*-mer frequency is not sufficient. In this study, we propose the “genomic style” inspired by deep-learning for image classification as yet another new sequence feature that replaces the *k*-mer frequency basically used in the existing binning methods.

In the field of deep learning-based image processing (Jing et al., 2019), image classification is largely divided into those based on shape and those based on style. The former is a classification based on the shape of what is drawn in the image, and the latter is a classification based on artistic style of an image such as color usage and texture, which is unique to the painter. Since the artistic style of an image is a rich descriptor that captures both visual and historical information about the painting, several studies have addressed the problem of style recognition employing the machine learning technique to automatically detect the artistic style of a painting (Lecoutre et al., 2017).

Recently, a deep learning approach to style transfer of images, Neural style transfer (Gatys et al., 2016) has been reported that generates an image converted to another style while maintaining the content (shape and arrangement of objects) in the image. The style matrix concept has been introduced as a method of expressing the artistic style. The style matrix consists of the correlation values between the feature maps of different filter responses in the same hidden layer of the convolutional neural network. This definition of style matrix has been hypothesized to be able to represent the style of the image, such as color usage and texture. One distinctive feature of the style matrix is that the style transfer task needs to transfer the style to a new content while preserving the content of an image, thus separating the style from the content as much as possible. This distinguished feature is very suitable for our purpose to extract the style of genome sequence specific to bacterial species while preserving the content of genetic information.

In this study, we hypothesized that the DNA sequence must have a style, called “genomic style” equivalent to the style of an image, which is unique to the bacterial species. We applied the style matrix concept developed in Neural style transfer to DNA sequences and aimed to extract genomic styles. Then, we proposed a method of clustering the metagenomic sequences using the extracted genomic style as sequence feature. We expect that the genomic style opens up new insights into the biological sequence analysis in metagenome analysis and bacterial taxonomy analysis.

## Methods

The outline of the method is as follows. First, we constructed a convolutional neural network (CNN) for modeling the content of DNA sequences, called the Master CNN model, and learned the species classification using known bacterial genome sequences. Second, we calculated the style matrix by inputting the DNA sequence into the trained master CNN model and extracted the style of DNA sequences as genomic style. Third, we performed binning by clustering DNA sequences using the obtained genomic style as a feature value. The source code for the implementation of this genomic style method, along with the dataset for the performance evaluation, is available at https://github.com/friendflower94/binning-style.

### Style of Image

First, we briefly introduce Neural Style Transfer (Gatys et al., 2016), a deep learning-based image-style transfer method that has been recently proposed to generate an image converted to another style while maintaining the content in the image. An image whose object layout is to be preserved is called a content image, and an image with a style used for conversion is called a style image. The style such as color and texture of the content image are rewritten while maintaining the position and outline of the object in the content image. To accomplish the style transfer task for an image, Neural style transfer proposed a method, called “style matrix” to express the style of an image by making use of VGG (Simonyan et al., 2014), a CNN specially designed for image recognition.

### Convolutional and Pooling Layer in CNN

The CNN, which has demonstrated an epoch-making performance in image analysis, has a convolution layer consisting of filters that automatically learns features of images such as straight lines and circles existing in the image. By combining this convolutional mechanism in multiple layers, deeper hierarchies acquire higher-order features such as facial contours. Another technical feature of CNN is that one image is learned to be decomposed into multiple components called *feature maps*.

More concretely, in the first convolutional layer, the inner product between the filter of arbitrary size and each local region of the input image is calculated. The filter scans the entire input image with some fixed stride. This process extracts the feature map of the input image. In the deeper hidden layer, the inner product between the filter and the feature map of the previous layer is calculated. Each hidden layer has multiple filters so that the hidden layer outputs the set of feature maps (output one feature map per one filter). The pooling layer calculates the maximum value or average value within a certain range and makes it the representative value within the range. This makes it possible to acquire robustness against the position movement and compress information in the input image. Also, the input to the convolutional layer may include multiple input vectors, called channels (which are also considered as feature maps in the input layer). For example, when the input to the CNN is an RGB color image, there are three channels: red (R), green (G), and blue (B).

For a filter function in the *l*-th hidden layer of the CNN, the input is the set of feature maps in the (*l-*1)-th hidden layer 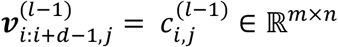, where *d* is the size of the filter, *m* is the size of feature map, and *n* is the number of feature maps. The output for the *k*-th filter is a feature map of the *l*-th layer 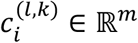, which is defined as follows:

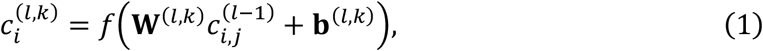

where *f* is an activation function, **W**^(*l,k*)^ ∈ ℝ^*m*×*n*×*d*^ is the weight matrix of the *k*-th filter in the *l*-th convolutional layer, and **b**^(*l,k*)^ is the bias vector.

### Style Matrix for Image Style

Neural Style Transfer (Gatys et al., 2016) proposed a style matrix as a method to represent the artistic style of an image. The style matrix consists of the correlation values between the multiple feature maps of different filter responses in the same hidden layer in VGG (also defined as the Gram matrix of feature maps). In their style transfer experiment to convert an image to another style while maintaining the content in the image, it was revealed that it was possible to separate the content information such as the shape and spatial position of the objects in the image from the style information such as color and texture by using the style matrix.

Neural style transfer used VGG (Simonyan et al., 2014) as an image recognition model. VGG16 is a CNN model composed of 13 convolutional layers, 5 pooling layers, and 3 fully connected layers. VGG obtained the second place in 2014 at ILSVRC, an image recognition competition. VGG is a model that has been trained by ImageNet (Krizhevsk et al., 2012), a dataset of over 14 million RGB images.

### Genomic Style

In this study, by applying the style matrix method of extracting the style to genome sequences, we extract genomic styles specific to bacterial species such as base composition and codon usage.

First, we constructed the master CNN model to perform bacterial species classification for modeling the content of DNA sequences. This master model is a CNN model in which eight modules consisting of the convolutional layer, batch normalization, rectified linear unit (ReLU) and the pooling layer are stacked, and finally, the output is obtained through the global average pooling layer and the fully connected layer (displayed in Figure 1). The cross-entropy error function for multi-class classification was used for the loss function to be minimized in learning. The input is a one-hot coding representation (one-hot vector) of four DNA nucleotides in a height of four dimensions and a width of 1,024 dimensions with the maximum length of DNA sequences. Several previous studies (Alipanahi *et al*., 2015; Zhou and Troyanskaya, 2015; Kelley *et al*., 2016; Zeng *et al*., 2016; Aoki and Sakakibara, 2018) have shown that a CNN can be applied to extraction of a sequence motif specifically conserved among target DNA sequences. When one-hot coding representation of four DNA nucleotides is employed, then a filter with a one-dimensional convolution operation applied temporally over a sequence can be considered a position weight matrix for representing a motif. Here, a “one-dimensional” convolution operation for sequences is interpreted as scanning the input sequence only in one direction along the sequence.

**Figure 1.**
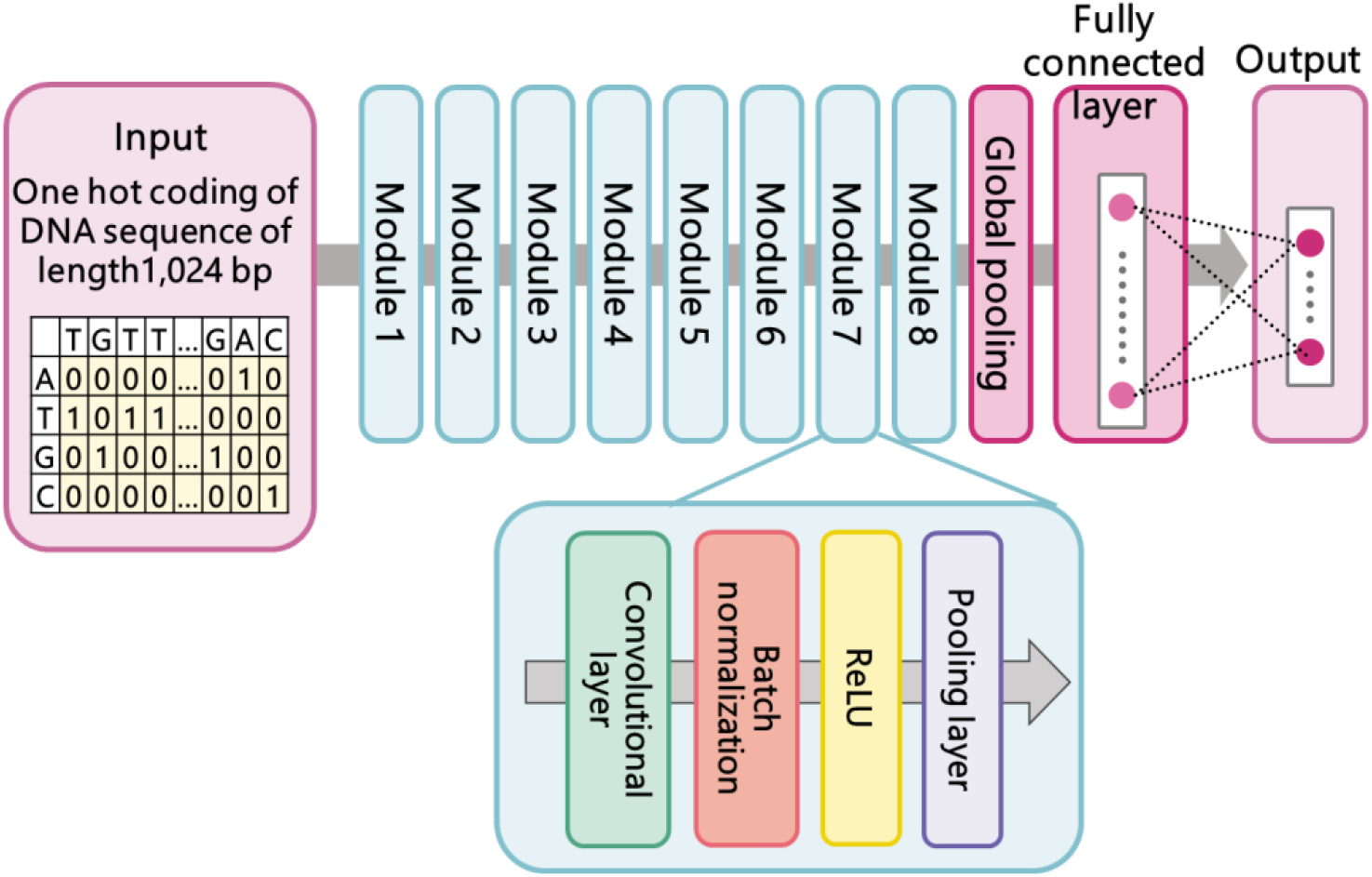
Master CNN model for modeling the content of DNA sequences. The master CNN model is trained for the task of bacterial species classification.

Second, the style matrix of the DNA sequence, called *genomic style*, was calculated using the master CNN model with the learned parameters after the species classification learning was completed. In the *l*-th hidden layer, when *N*_*l*_ denotes the number of feature maps and *M*_*l*_ denotes the size of the feature map, the *l*-th layer output *F*^(*l*)^ becomes 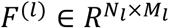. Let 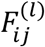 denote the *j*-th element of the *i*-th feature map in the *l*-th layer. Then, the element 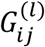 of the style matrix *G*^(*l*)^of the *l*-th layer is defined to be the inner product between the *i*-th feature map and the *j*-th feature map in the *l*-th layer, as formulated in the following formula and displayed in Figure 2:

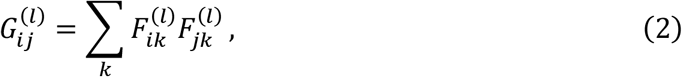

By inputting the DNA sequence into the master CNN model, the style matrix of the input DNA sequence is calculated by the above formula. Then, we use this style matrix as the feature vector for clustering the DNA sequences.

**Figure 2.**
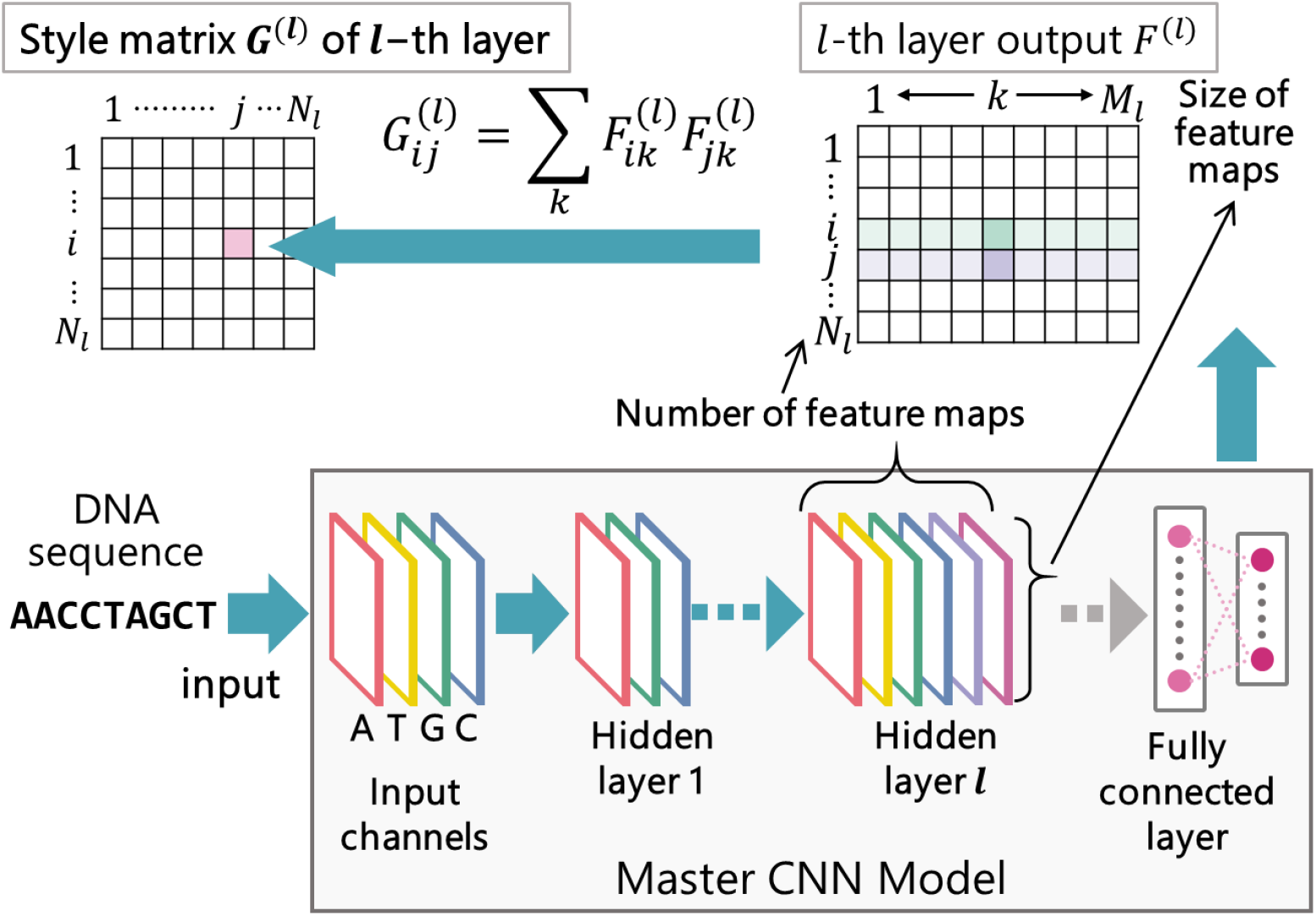
Calculation of style matrix 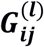 of the *l*-th hidden layer. The style matrix of the DNA sequence, called *genomic style*, is calculated using the master CNN model for the input DNA sequence.

### Binning using Genomic Style

Binning is the process of clustering DNA sequences into bins of the same species in a mixed sample of DNA sequences from multiple species (Thomas et al., 2012). We perform binning by clustering DNA sequences using the obtained style matrix as a feature vector. Since the style matrix *G*^(*l*)^ can be obtained for each layer, the style matrix is calculated for the first to sixth layers, that is, *l* = 1, 2, 3, 4, 5, and 6, and binning is performed using each of them in the following experiment.

The input to the master CNN model has a fixed sequence length of 1,024 bp. Since the length of the DNA sequence to be binned is longer than 1,024 bp, the input sequence is scanned with a fixed length of 1,024 bp and shifted by 500 bp, and the style matrix for each 1,024 bp is calculated. Then, the average of them is used as the style matrix for the entire input sequence. We employed the Agglomerative Clustering method (Frigui et al., 1997) for clustering with the style matrix. Agglomerative clustering is a method that starts with a cluster consisting of one element and recursively merges two clusters with the minimum inter-cluster distance until the target number of clusters is reached. The Euclidean distance of the style matrix was used for the distance between elements, and the distance between clusters is calculated using the Ward method.

In addition, in order to compare with state-of-the-art binning tools, we incorporated the coverage (abundance) information into binning by the style matrix. The extension simply added another dimension into the feature vector to represent the coverage value of each DNA sequence. We adjusted the scale of the coverage value to fit the scale of the style matrix. Agglomerative clustering with Euclidean distance was applied to the extended feature vector. We call this extension “style matrix with coverage”.

### Existing Binning Methods for Performance Comparison

The purpose of our experiment is to compare the ability of the proposed sequence feature, genomic style in replace of *k*-mer frequency for the binning task.

As the baseline for performance comparison, binning was performed using “*k*-mers frequency”. For each DNA sequence, the number of occurrences of sub-sequences of length *k* was counted by scanning and shifting by one base. The number of occurrences was divided by the total number to calculate the *k*-mers frequency. The *k*-mers frequency was calculated for *k*=3 and 4, and clustering was performed using an agglomerative clustering with *k*-mers frequency as the feature vector. In addition, MetaBAT2 (Kang et al., 2019) has an option to process binning without abundance information and only use the tetra-nucleotide frequency (TNF). Therefore, MetaBAT2 without option of abundance scores was included in this performance comparison. MetaBAT (Kang et al., 2015) is an automated binning software that can process large datasets of DNA sequences. It is based on empirical probabilistic distances of TNF, that is, 4-mers frequency and genome abundance. MetaBAT is widely used by the microbiology community, and a new version of MetaBAT has been released, MetaBAT2 (Kang et al., 2019), which is a state-of-the-art binning tool and automates tuning of the parameter to obtain better binning results.

To compare the performance with state-of-the-art binning tools, the following tools were applied in our experiment: MetaBAT2 (with option to use abundance scores), CONCOCT (Alneberg et al., 2014), MaxBin2 (Wu et al., 2015), and MrGBP (Kouchaki et al, 2019). Default values were used for all parameters in these methods. MetaProb (Girotto et al, 2016) failed to accomplish the binning experiment on CAMI challenge dataset, therefore we removed it from our experiment. Note that all of these existing binning tools utilize coverage (abundance) information for each sequence as well as *k*-mers frequency.

### Datasets

The experiment proceeded with four different datasets. The first dataset was used for training the master CNN model. The second one consisted of assembled contigs to compare the effectiveness of genomic style and *k*-mer frequency as sequence features for the binning task, and the third one was obtained from Critical Assessment of Metagenome Interpretation (CAMI) challenge dataset (Sczyrba et al., 2017) for the performance comparison with the existing binning methods. The fourth one comprised random DNA sequences with different GC content.

### Training Dataset

As the training set, DNA sequences of bacterial whole genomes obtained from the National Center for Biotechnology Information (NCBI) Nucleotide database (Sayers et al., 2019) were used. There were 206 different taxa represented at the species level. The dataset was divided into 139 species and 92 species, allowing for 25 duplicates, and 139 species as training data to train the model. To make the dataset for training the master CNN model, we used 139 whole genomes, each one belonging to different species. Then, DNA sequences of 1,024 bp in length were randomly extracted from each genome. As a result, we constructed 128,000 DNA sequences as the training dataset.

### Binning Test Dataset to Compare Genomic Style and K-mer Frequency

92 species were used for the binning test data to compare genomic style and *k*-mer frequency. To produce the binning test dataset for assembled contigs, we used the software CAMISIM (Fritz et al., 2019) to simulate metagenomes. We used the ART read simulator to generate reads from a set of pre-specified genomes. It produced reads simulated from the Illumina HiSeq150 sequencing technology. The fragment size mean was set to 270 bp, and the fragment standard deviation was set to 27 bp. We used the golden standard assembly generated by CAMISIM, i.e., the set of contigs with their associated ground-truth labels. Then, for each contig, we kept contigs with a length over 10,000 bp and cropped the larger ones to obtain contigs of length 10,000 bp. We proceeded seven times to finally get a binning test dataset consisting of 86,346 contigs, each of length 10,000 bp and originating from 92 different species. For a distribution of taxa among the binning dataset, 67 species were part of the binning set but not of the training set; these species served as “unknown species” that underwent binning.

In addition, binning test datasets were created at the genus and family levels, respectively. In the genus-level test dataset, one species from each of the 30 genera was selected, and 12 species were not included in the training set. In the family-level test dataset, one species from each of the 16 families was selected, and 5 species were not included in the training set. In these two datasets, the binning task is expected to be easier and its classification accuracy will be higher.

### CAMI Challenge Dataset

The CAMI challenge (Sayers et al., 2019) is an effort to provide gold standard benchmark datasets for comparing tools for the analysis of metagenomic data. The reads in the low-complexity dataset of CAMI challenge are simulated with Illumina HiSeq error profile from 40 genomes and 20 circular elements. The gold-standard metagenome assembly constructed from the reads in the CAMI low-complexity dataset contains 19,499 DNA sequences (a set of contigs with their associated ground-truth labels), which is mixtures of variable-length fragments originating from individual species. This DNA sequence dataset is provided as the binning challenge for genome binning (unsupervised binning, that is clustering) to group sequences into unlabeled bins and taxonomic binning (supervised binning) to group the sequences into bins with a taxonomic label attached. Figure 3 also shows, for each taxonomic rank, the number of taxa which are exclusive to our training dataset, which are exclusive to the CAMI low-complexity dataset, and which are common to both. Note that the 40 genomes and 20 circular elements in the low-complexity dataset included those that were unidentified and those identified up to higher taxonomy in each taxonomic rank.

**Figure 3.**
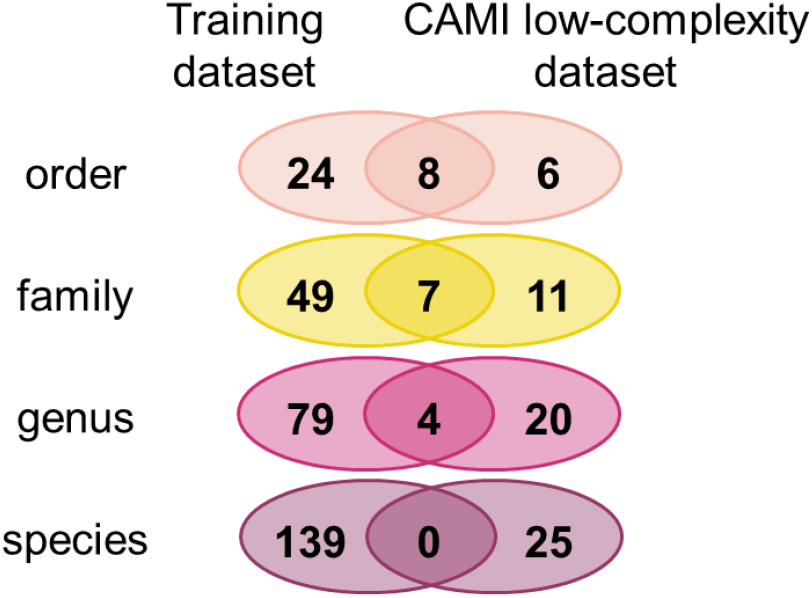
Common and different taxa between the training dataset and the CAMI low-complexity dataset. For each taxonomic rank, the number of taxa which are exclusive to the training dataset, which are exclusive to the CAMI low-complexity dataset, and which are common to both are shown. Note that the 40 genomes and 20 circular elements in the low-complexity dataset included those that were unidentified and those identified up to higher taxonomy in each taxonomic rank.

### Random DNA-Sequences with Different GC-Content

Random DNA sequences with different GC content were generated to evaluate what sequence features the genome-style matrix has acquired. 500 random DNA sequences were generated for each of two groups with different GC contents of x% and (100−x)%. For a total of 1,000 generated sequences, we calculated the style matrix and performed clustering by agglomerative clustering. The binning performance was evaluated with GC content of x=53, 55, 57.

## Results

The following measures were used for the performance evaluation criteria: Adjusted Rand Index (ARI), Homogeneity and Completeness metrics.

The ARI (Hubert et al., 1985) is a metric derived from the Rand Index (RI) defined as:

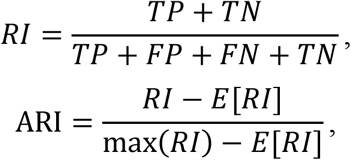

where *TP* denotes the number of the same species DNA-sequences in the same cluster, *TN* denotes the number of different species sequences in different clusters, *FP* denotes the number of different species sequences in the same cluster, and *FN* denotes the number of different species sequences in the same cluster. In many cases, the number of sequence pairs in different clusters is greater than the number of sequence pairs in the same cluster, so the naive clustering to divide all sequences into different clusters will increase the RI value. Therefore, ARI is obtained by decreasing as a penalty the value of RI when clustering is performed arbitrarily. Homogeneity is an index that measures the purity of bins without contamination. In the predicted cluster, if only elements from the same species belong, it is defined to be homogeneity. Completeness is a measure of whether elements belonging to the same species are assigned to the same cluster. Generally, the higher the homogeneity, the lower the completeness is, and the higher the completeness, the lower the homogeneity. Thus, homogeneity and completeness are trade-off indices.

### Effectiveness of Genomic Style as Sequence Features for Binning Task

Table 1 shows the binning accuracy using the style matrix from the 1st to 6th layer, the *k*-mer frequency (*k*=3, 4) and MetaBAT2 without option of abundance scores for the binning test datasets at the species level, genus level and family level. Among the style matrices using 6 different layers, the 4th layer gave the highest accuracy in all three accuracy indices in the species and family-level test datasets. In the genus-level test dataset, the 5th layer showed the highest accuracy in ARI and completeness scores, but the differences between 4th and 5th layers were small. When comparing among three test datasets at the species, genus, and family levels, the higher the taxonomic level, the higher the accuracy. In particular, the ARI score using 4th layer reached 0.890 for the family-level test dataset.

**Table 1.** Performance evaluation of the binning methods using the style matrix from the 1st to 6th layer, the *k*-mer frequency (*k*=3, 4), and MetaBAT2 (without option of abundance scores). Three accuracy indices, ARI, homogeneity, and completeness are shown for the binning test datasets at the species level, genus level and family level.

| Method | Style matrix (of layer $l$) | | | | | | $k$-mers frequency | | MetaBAT2 |
| --- | --- | --- | --- | --- | --- | --- | --- | --- | --- |
| | $l = 1$ | $l = 2$ | $l = 3$ | $l = 4$ | $l = 5$ | $l = 6$ | $k = 3$ | $k = 4$ | |
| Species |  |  |  |  |  |  |  |  |  |
| ARI | 0.316 | 0.346 | 0.377 | **0.441** | 0.410 | 0.290 | 0.292 | 0.308 | 0.349 |
| Homogeneity | 0.654 | 0.681 | 0.704 | 0.714 | 0.667 | 0.589 | 0.625 | 0.650 | **0.835** |
| Completeness | 0.628 | 0.656 | 0.682 | **0.707** | 0.689 | 0.641 | 0.596 | 0.621 | 0.588 |
| Genus |  |  |  |  |  |  |  |  |  |
| ARI | 0.458 | 0.450 | 0.530 | 0.571 | **0.604** | 0.517 | 0.427 | 0.438 | 0.436 |
| Homogeneity | 0.713 | 0.733 | 0.758 | 0.765 | 0.747 | 0.690 | 0.706 | 0.733 | **0.930** |
| Completeness | 0.679 | 0.695 | 0.735 | 0.760 | **0.763** | 0.715 | 0.654 | 0.686 | 0.533 |
| Family |  |  |  |  |  |  |  |  |  |
| ARI | 0.621 | 0.748 | 0.693 | **0.890** | 0.838 | 0.811 | 0.649 | 0.622 | 0.505 |
| Homogeneity | 0.796 | 0.879 | 0.850 | 0.900 | 0.865 | 0.841 | 0.799 | 0.798 | **0.991** |
| Completeness | 0.745 | 0.817 | 0.790 | **0.869** | 0.840 | 0.819 | 0.739 | 0.732 | 0.518 |

Compared with the *k*-mer frequency and MetaBAT2 (without option of abundance scores), at all taxonomic levels, the accuracies using the style matrix were highest for ARI and completeness while MetaBAT2 exhibited the highest homogeneity. Since MetaBAT2 employs a variant of the 4-mers frequency (TNF) as the sequence feature, the difference of binning accuracy between the 4-mers frequency and MetaBAT2 depends on their clustering methods; MetaBAT2 used a graph clustering method, whereas binning based on 4-mers frequency used the agglomerative clustering.

### Performance Comparison with Existing Binning Tools

The binning performance of the style matrix with coverage was compared with that of state-of-the-art binning tools on the CAMI challenge dataset. The style matrix with coverage used 4th layer, which exhibited the highest performance in the binning test without using the coverage score. The result is shown in Tables 2.

**Table 2.** Performance comparison with existing binning methods on CAMI challenge dataset.

| | Style matrix with coverage ($l = 4$) | Style matrix ($l = 4$) | MetaBAT2 | CONCOCT | MaxBin2 | MrGBP |
| --- | --- | --- | --- | --- | --- | --- |
| ARI | **0.207** | 0.160 | 0.044 | 0.165 | 0.146 | 0.000 |
| Homogeneity | 0.646 | 0.563 | **0.979** | 0.925 | 0.894 | 0.570 |
| Completeness | **0.400** | 0.335 | 0.243 | 0.282 | 0.270 | 0.195 |
| Num. of outliers | 0 | 0 | 15,592 | 11,063 | 11,182 | 11,091 |

Style matrix of 4th layer with coverage achieved the highest ARI and completeness scores among all binning methods while MetaBAT2, CONCOCT and MaxBin2 showed very high homogeneity. The performance difference between the style matrix of 4th layer with coverage and CONCOCT (which exhibited the highest ARI among the existing tools) was significant in the ARI score. All existing tools excluded more than half of DNA sequences in CAMI challenge dataset as outliers, which resulted in very low ARI and completeness scores. Comparing the style matrix of 4th layer without coverage information, it was revealed that coverage information helped improve accuracy of ARI, homogeneity and completeness scores.

### Binning DNA-Sequences with Different GC-Content

Table 3 shows the result of clustering 1,000 randomly generated sequences with different GC content by using the style matrix of 2nd and 4th layer. Figure 4 displays the visualization of clustering the DNA sequences by reducing the dimension of the style matrix to two dimensions using UMAP. UMAP (McInnes et al., 2018) is a state-of-the-art dimension compression tool that performs dimension reduction considering nonlinear components at high speed. It turned out that both style matrices of 2nd and 4th layers could separate sequences with different GC content, while the style matrix of 2nd layer showed higher accuracy than the 4th layer. In the plots displayed in Figure 6, DNA sequences with larger difference in GC content were more clearly separated, and the style matrix of 2nd layer separated the two groups with different GC content more explicitly than the style matrix of 4th layer.

**Table 3.** Binning accuracy using the style matrix of 2nd and 4th for random DNA sequences with different GC content.

| Method | ARI | Homogeneity | Completeness |
| --- | --- | --- | --- |
| Group 1 (GC:53%), Group 2 (GC:47%) |  |  |  |
| Style matrix of layer 2 | 0.732 | 0.627 | 0.627 |
| Style matrix of layer 4 | 0.000 | 0.000 | 0.000 |
| Group 1 (GC:55%), Group 2 (GC:55%) |  |  |  |
| Style matrix of layer 2 | 1.000 | 1.000 | 1.000 |
| Style matrix of layer 4 | 0.611 | 0.503 | 0.504 |
| Group 1 (GC:57%), Group 2 (GC:43%) |  |  |  |
| Style matrix of layer 2 | 1.000 | 1.000 | 1.000 |
| Style matrix of layer 4 | 0.910 | 0.847 | 0.848 |

**Figure 4.**
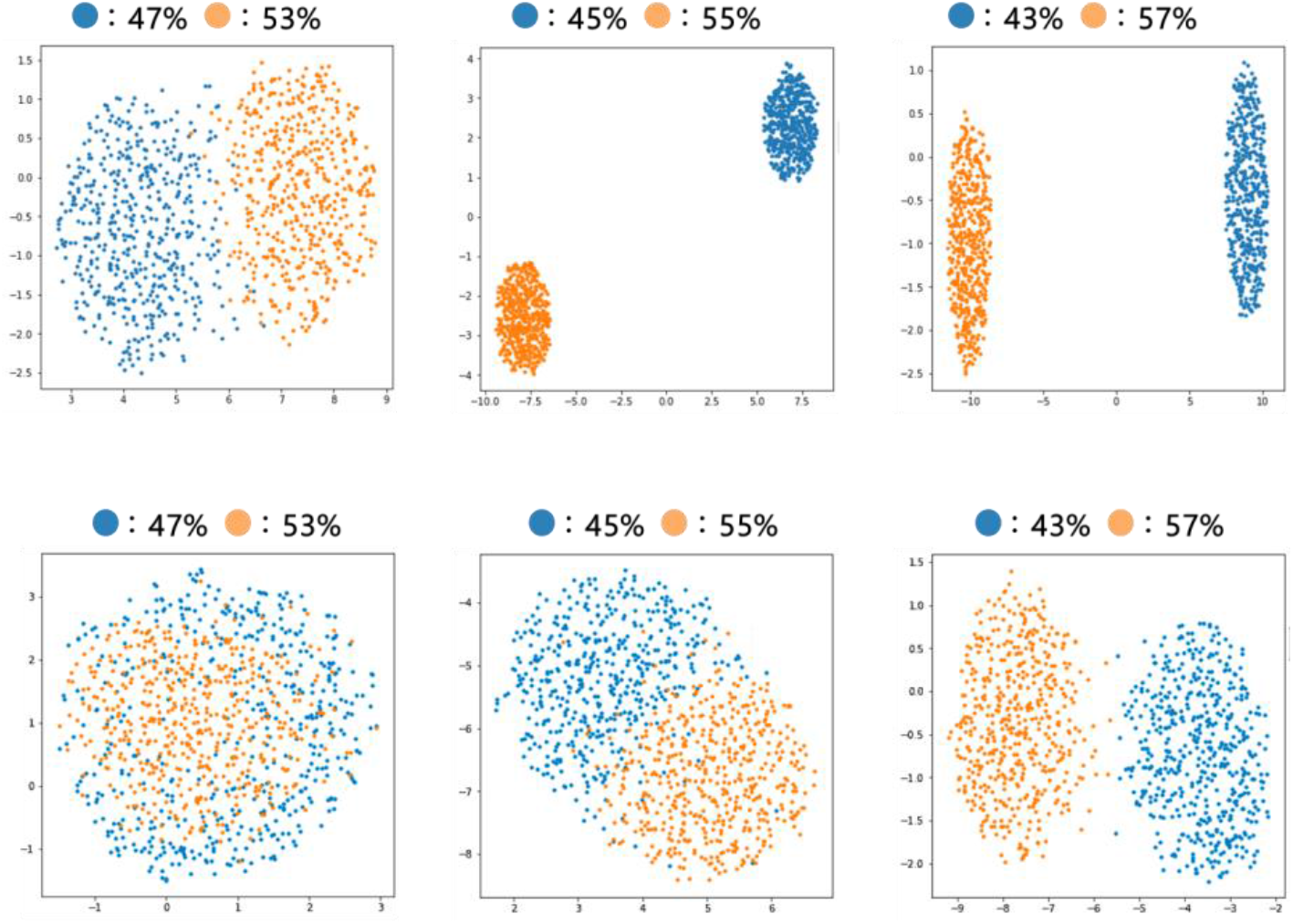
Visualization of the results of clustering DNA sequences with different GC content by reducing the dimension of the style matrix to two dimensions using UMAP. (Upper) Clustering using the style matrix of 2nd layer. (Lower) Clustering using the style matrix of 4th layer.

## Discussion

The accuracy of binning with the style matrix of the master CNN model constructed in this study was the highest when the style matrix of 4th layer was used. In the performance comparison with existing binning tools, the style matrix of 4th layer with coverage achieved the highest accuracy in ARI and completeness compared to state-of-the-art methods. With regard to homogeneity, MetaBAT2, CONCOCT and MaxBin2 showed higher accuracy than other methods. These methods excluded more than half of DNA sequences in CAMI challenge dataset as outliers. The outliers form bins of only one element. Homogeneity measures whether each cluster contains only elements from the same species. Obviously, bins containing one element are homogeneous, and the homogeneity value tends to be higher when generating more bins with a smaller number of elements.

The evaluation of the style matrix for the generated sequence with different GC content revealed that the style matrix of both 2nd layer and 4th layer could distinguish the difference in the GC content. Therefore, it can be said that the style matrix looks at differences in GC content and base composition depending on the bacterial species. In the 2nd layer, the sequence can be separated to some extent even if the GC content difference is 6%, but in the 4th layer, the sequence cannot be separated unless the difference exceeds 10% in GC content. Therefore, it can be considered that the GC content is captured in the shallow layer.

The feature map itself in the master CNN model did not work as a feature for DNA sequence clustering (unsupervised classification). The feature map is the filter responses from the previous layer, that is, the inner product between the filter and each local region of the previous layer’s output (this is also feature map), as defined in the formula (1). The feature map could acquire some features for the content of the input DNA-sequence. However, Table 4 shows that the binning accuracies using feature maps of the 2nd, 4th and 6th layers were very low. This result indicated that the calculation of the “correlation values” (that is, the style matrix) between multiple feature maps was necessary as the sequence feature for binning DNA sequences.

**Table 4.** Binning accuracy using the feature maps as sequence features for clustering. The accuracy is shown for using feature maps of 2nd, 4th and 6th layers.

| Method | ARI | Homogeneity | Completeness |
| --- | --- | --- | --- |
| Feature map of layer 2 | 0.010 | 0.074 | 0.071 |
| Feature map of layer 4 | 0.003 | 0.034 | 0.032 |
| Feature map of layer 6 | 0.002 | 0.029 | 0.027 |

As stated in the introduction section, when one-hot coding representation of four DNA nucleotides is used in the CNN, then a filter with a one-dimensional convolution operation can be considered a position weight matrix representing a motif (Alipanahi *et al*., 2015; Zeng *et al*., 2016; Aoki and Sakakibara, 2018). The position weight matrix is a generalization of *k*-mer (word of length *k*). Since our method using the style matrix of the *l*-th layer scans the DNA sequence with filters of the *l*-th layer and counts the matches with the position weight matrix represented by the filter, the style matrix can be considered as a generalization of the *k*-mers frequency.

More precisely, in the process of pre-training the master CNN as the first step of binning, sequence and structure motifs were extracted within the filters that were important for taxonomic classification. The feature map is the filter responses for the input DNA sequence and the genomic style is defined as the gram matrix of feature maps, hence it represents the preference of those motifs. The filters in the master CNN could be considered position weight matrix representing motifs, hence the gram matrix of feature maps could be considered the preference distribution of probabilistic sequence motifs, include the concept of probabilistic sequence signature (MetaProb), probabilistic *k*-mers statistics (MetaCon), empirical probabilistic distances of TNF (MetaBAT2), textual representations of sequence data (MrGBP), the distribution of a carefully selected set of *k*-mers (MetaCluster, (Yang et al., 2010)). Therefore, the genomic style could offer the unified framework for these variations of *k*-mers frequency.

## Conclusions

In this study, we proposed a new concept called “genomic style”, which is a feature of genomic sequences that are not limited to base composition and *k*-mers frequency, as well as a method for binning metagenomic sequences using the genomic style. We first constructed a CNN-based master model for modeling the content of DNA sequence and performed species classification learning. Genomic style was extracted from the style matrix calculated using the trained master CNN model, and binning of the metagenome sequence was performed using the style matrix. The binning results revealed that the genomic style could be a DNA sequence feature unique to the bacterial species for the accurate DNA sequence classification. We also investigated what kind of sequence features the style matrix acquired. It turned out that some basic sequence features such as GC content and DNA motifs were captured in the style matrix of shallow layers.

In some previous studies (Simonyan et al., 2014) using the CNN for image recognition, it was revealed that the deeper hidden layers captured higher-order features of the image. It remains as a future work to clarify what kind of higher-order features of DNA sequence could be captured by the style matrix in a deeper layer. In order to improve the accuracy of binning of the metagenome, it is also our future work to improve the master CNN model to extract more accurate genomic styles specific to the bacterial species.

## List of Abbreviations

CNN: Convolutional Neural Network
TNF: Tetra-Nucleotide Frequency
ARI: Adjusted Rand Index

## Availability

The source code for the implementation of this genomic style method, along with the dataset for the performance evaluation, is available at https://github.com/friendflower94/binning-style.

## Declarations

### Authors’ Contributions

YY; implemented the software, analysed data, and compared with the existing methods. AH; implemented the software and analysed data. AY, MA; analysed data. YS; designed and supervised the research, analysed data, and wrote the paper. All authors read and approved the final manuscript.

## References

Alipanahi, B. et al. (2015) Predicting the sequence specificities of DNA- and RNA-binding proteins by deep learning. Nat. Biotechnol., 33, 831–838.

Alneberg, J. et al. (2014) Binning metagenomic contigs by coverage and composition. Nat. Methods, 11(11), 1144–1146, 1063-1071.

Aoki, G. and Sakakibara, Y. (2018) Convolutional neural networks for classification of alignments of non-coding RNA. Bioinformatics, 34(13), i237–i244.

Bishop, CM. (2006) Pattern Recognition and Machine Learning. Springer-Verlag, Berlin, Heidelberg.

Chatterji, S. et al. (2008) Compostbin: a DNA composition-based algorithm for binning environmental shotgun reads. Res. Comput. Mol. Biol., 17–28.

Durbin, R. et al. (1998) Biological Sequence Analysis: Probabilistic Models of Proteins and Nucleic Acids. Cambridge University Press, Cambridge, UK.

Frigui, H., & Krishnapuram, R. (1997). Clustering by competitive agglomeration. Pattern. Recognit., 30(7), 1109–1119.

Fritz, A. et al. (2019) CAMISIM: simulating metagenomes and microbial communities. Microbiome, 7(1), 17.

Gatys, L.A. et al. (2015) A neural algorithm of artistic style. arXiv preprint, 1508.06576.

Girotto, S., Pizzi, C., Comin, M. (2016) MetaProb: accurate metagenomic reads binning based on probabilistic sequence signatures. Bioinformatics, 32(17), i567–i575.

Huson, D.H. et al. (2007) MEGAN analysis of metagenomic data. Genome Res., 17, 377–386.

Hubert, L. and Arabie, P. (1985) Comparing partitions. J. Classif., 2(1), 193–218.

Jing, Y. et al. (2019) Neural style transfer: A review. IEEE Trans. Vis. Comput. Graph, doi: 10.1109/TVCG.2019.2921336.

Kang, D. et al. (2019) MetaBAT 2: an adaptive binning algorithm for robust and efficient genome reconstruction from metagenome assemblies. PeerJ, 7, e7359.

Kelley, D.R. et al. (2016) Basset: learning the regulatory code of the accessible genome with deep convolutional neural networks. Genome. Res., 26, 990–999.

Kouchaki., S., Tapinos., A., Robertson, D. L. (2019) A signal processing method for alignment-free metagenomic binning: multi-resolution genomic binary patterns. Sci. Rep., 9(1), 2159.

Krizhevsky, A. et al. (2012) ImageNet classification with deep convolutional neural networks. In Advances in Neural Information Processing Systems, pages 1097–1105.

Lu, Y.Y. et al. (2017) COCACOLA: binning metagenomic contigs using sequence composition, read coverage, co-alignment and paired-end read linkage. Bioinformatics, 33(6):791–798.

Mande, S.S., Mohammed, M.H. and Ghosh, T.S. (2012) Classification of metagenomic sequences: methods and challenges. Brief. Bioinform., 13(6), 669–681.

McInnes, L. et al. (2018) UMAP: Uniform manifold approximation and projection. J. Open. Source. Softw., 3(29), 861.

Sayers, E.W. et al. (2019) Genbank. Nucleic. Acids. Res., 47(D1), D94–D99. (National Center for Biotechnology Information, U.S. National Library of Medicine, https://www.ncbi.nlm.nih.gov/nucleotide/)

Sczyrba, A. et al. (2017) Critical Assessment of Metagnome Interpretation -a benchmark of metagenomics software. Nat. Methods, 14, 1063–1071.

Simonyan, K. and Zisserman, A. (2014) Very deep convolutional networks for large-scale image recognition. arXiv preprint, 1409.1556.

Tareen, A. and Kinney, J.B. (2019) Logomaker: beautiful sequence logos in Python. BioRxiv, 635029.

Thomas, T. et al. (2012). Metagenomics - a guide from sampling to data analysis. Microb. Inform. Exp., 2(1), 3.

Wood, D.E. and Salzberg, S.L. (2014) Kraken: ultrafast metagenomic sequence classification using exact alignments. Genome Biol., 15, R46.

Wu, Y-W., Simmons, B.A., Singer, S.W. (2015). Maxbin 2.0: an automated binning algorithm to recover genomes from multiple metagenomic datasets. Bioinformatics, 32(4), 605–607.

Yang, B. et al. (2010) Unsupervised binning of environmental genomic fragments based on an error robust selection of l-mers. BMC Bioinformatics, 11, S5.

Zeng, H. et al. (2016) Convolutional neural network architectures for predicting DNA-protein binding. Bioinformatics, 32, i121–i127.

Zhou, J. and Troyanskaya, O.G. (2015) Predicting effects of noncoding variants with deep learning-based sequence model. Nat. Methods., 12, 931–934.

